# Deamidation in Moxetumomab Pasudotox Leading to Conformational Change and Immunotoxin Activity Loss

**DOI:** 10.1101/761635

**Authors:** X. Lu, S. Lin, N. De Mel, A. Parupudi, M. Pandey, J. Delmar, X. Wang, J. Wang

## Abstract

Asparagine deamidation is a common posttranslational modification in which asparagine is converted to aspartic acid or isoaspartic acid. By introducing a negative charge, deamidation could potentially impact the binding interface and biological activities of protein therapeutics. We identified a deamidation variant in moxetumomab pasudotox, an immunotoxin Fv fusion protein drug derived from a 38-kilodalton (kDa) truncated *Pseudomonas* exotoxin A (PE38) for the treatment of hairy-cell leukemia. Although the deamidation site, Asn-358, was outside of the binding interface, the modification had a significant impact on the biological activity of moxetumomab pasudotox. Surprisingly, the variant eluted earlier than its unmodified form on anion exchange chromatography, which suggests a higher positive charge. Here we describe the characterization of the deamidation variant with differential scanning calorimetry and hydrogen-deuterium exchange mass spectrometry, which revealed that the Asn-358 deamidation caused the conformational changes in the catalytic domain of the PE38 region. These results provide a possible explanation for why the deamidation affected the biological activity of moxetumomab pasudotox and suggest an approach that can be used for process control to ensure product quality and process consistency.

**Statement of Significance:** Asparagine deamidation can have a potentially significant impact on protein therapeutics. Previous studies have revealed that deamidation at a single site significantly reduces the biological activity of moxetumomab pasudotox, a recombinant anti-CD22 immunotoxin developed for the treatment of B-cell malignancies. Surprisingly, despite the fact that deamidation introduced a negative charge, the deamidated variant eluted earlier than its unmodified form on anion exchange chromatography. In order to understand these observations, we isolated the deamidated variant using an anion exchange column and characterized it by differential scanning calorimetry and hydrogen-deuterium exchange mass spectrometry. The results revealed the conformational change caused by the deamidation, which explains the diminished biological activity of the variant and its early elution time on anion exchange chromatography.

## INTRODUCTION

Since the approval of recombinant human insulin by the U.S. Food and Drug Administration (FDA) in 1982, the use of therapeutic proteins has grown tremendously for treatment of many diseases and disorders, including cancer, autoimmune and inflammatory disorders, and metabolic, cardiovascular, and infectious diseases (1, 2). Protein-based biopharmaceuticals often have higher specificity and therefore lower potential to cause adverse effects and immune responses than small-molecule drugs, but they also present their own challenges (2). One of the most critical of these challenges is the chemical degradation that may occur during the manufacturing and storage for most therapeutic proteins, which leads to one or more posttranslational modifications (PTMs) (2, 3). Asparagine (Asn) deamidation, one of most common nonenzymatic PTMs, has long been considered as a major degradation pathway for many monoclonal antibodies and other therapeutic proteins (4). It is generally believed that under neutral and basic conditions, the reaction goes through a five-membered cyclic succinimide intermediate, which then hydrolyzes and yields both isoaspartic acid (iso-Asp) and Aspartic acid (Asp) (5–7).

It has been reported that some PTMs could have a significant impact on the biological activity of therapeutic proteins. For example, in a study of dornase alpha, a recombinant enzyme used for the treatment of cystic fibrosis, deamidation of a single Asn residue resulted in loss of its biological activity (8). Deamidation was also reported to compromise antibody colloidal stability and to increase aggregation under low-pH conditions (9). Especially in light of the relationship between protein structure and function, detailed structural characterization is essential to understand the molecular basis of such reactions.

Hydrogen-deuterium exchange (HDX) mass spectrometry (MS) has emerged as a useful tool to monitor the conformation and dynamics of therapeutic proteins (10–13). Conventional biophysical techniques such as circular dichroism and fluorescence (14–16) provide only general information, and it is difficult to detect subtle conformational changes or changes related only to specific regions of proteins with these techniques. In contrast, HDX-MS can be used to detect and localize the impact of PTMs on the structure of proteins at the peptide level, or even at the residue-level. For example, HDX has been used to study Asp isomerization and methionine oxidation in complementarity-determining regions (CDR) of antibodies. Although they are only one amino acid apart in sequence, these two modifications have been found to result in opposing conformational impacts on local and nearby regions, leading directly to different alterations on antibody-antigen binding affinity (17). It has also been reported that CDR oxidation of tryptophan significantly reduces antigen binding affinity, and HDX-MS data have revealed that the modifications increased the flexibility of variable regions and disrupted local conformation (18). While a similar experiment with Asn deamidation revealed no measurable local conformational changes caused by the modification, this finding may have been compromised by the low deamidation levels (~30%) in the material used in the study (19).

A few studies have explored how Asn deamidation affects the 3-dimensional structure of proteins (20–25). Deamidation of human norovirus capsid protein has been reported to attenuate glycan binding, and HDX-MS data have revealed localized alterations in protein flexibility, supporting findings from nuclear magnetic resonance and X-ray crystallography studies (26). In another HDX-MS study, deamidation was reported to slow Curli amyloid-protein aggregation (11).

Moxetumomab pasudotox (formerly CAT-8015) is a recombinant immunotoxin developed for the treatment of B-cell malignancies (27) that received FDA approval in September 2018 for the treatment of adults with relapsed or refractory hairy-cell leukemia who have received at least 2 prior systemic therapies, including treatment with a purine nucleoside analog (28, 29). It is composed of a 38-kDa truncated *Pseudomonas* exotoxin A (PE38) that is genetically fused to the variable fragment (Fv) of an anti-CD22 antibody. When bound to CD22, a B-lymphocyte–specific cell surface protein, it is rapidly internalized and processed. The *Pseudomonas* exotoxin (PE) catalyzes the transfer of ADP ribose from nicotinamide adenine dinucleotide (NAD) to elongation factor 2 (eEF2), which inhibits protein translation and leads to cell death.

Deamidation in the catalytic domain of PE38 at Asn-358 has previously demonstrated significantly reduced its biological activity in cell-based bioassay (30), but it remains unclear how a single residue modification could have such a marked impact on the cytotoxicity of the molecule. Another puzzling observation is that the deamidated species eluted before the major product peak on a positively charged anion exchange column, despite the fact that Asn deamidation introduces an extra negative charge.

In this work, Asn-358–deamidated moxetumomab pasudotox was first isolated using ion exchange chromatography (IEC). The structural impact of deamidation was carefully examined by both differential scanning calorimetry (DSC) and HDX-MS, which showed alteration of the conformation of the PE catalytic domain. In the comprehensive functional studies described here, we gained a deep understanding of why the single deamidation diminished biological activity and appeared as a basic variant, an early elution peak in anion exchange chromatography. After evaluating the results from our stability study and the refolding process, a control strategy was established and implemented in the commercial process to minimize the deamidation level and maintain consistency.

## MATERIALS AND METHODS

### Moxetumomab pasudotox

Recombinant moxetumomab pasudotox used in this study was expressed in *Escherichia coli* strains and was purified at AstraZeneca (Gaithersburg, MD) as previously described (31). All other commercial chemicals were reagent grade or better. Peptide mapping and cell-based bioassay methods were employed as previously described (30).

### IEC fractionation

IEC fractionation was performed on a 1200 HPLC system equipped with a PL-SAX strong anion exchange column (4.6 × 50 mm, 8 μm, 1000 Å; Agilent, Santa Clara, CA). Mobile phase A was composed of 10 mM Tris and 1 mM ethylenediaminetetraacetic acid, pH 8.0, and mobile phase B was composed of 10 mM Tris, 1 mM ethylenediaminetetraacetic acid, and 0.5 M NaCl, pH 8.0. For each experiment, 500 μL of moxetumomab pasudotox starting material at a concentration of 1 mg/mL was loaded on the system and eluted with a linear gradient of up to 80% mobile phase B at a flow rate of 1.0 mL/min. The eluted protein was monitored by ultraviolet absorbance at 280 nm, and 2 fractions (pre-peak and main peak) were collected with a fraction collector. The fractions from multiple injections were pooled together and buffer exchanged with 25 mM sodium phosphate, pH 7.4, to a final concentration of approximately 1 mg/mL. The purity of IEC fractionation was verified by injecting 25 μg of starting material and 5 μg of each fraction on the anion exchange column.

### Differential scanning calorimetry

DSC experiments were performed with Microcal VP-Capillary DSC (Malvern Instruments, Malvern, PA) at a concentration of approximately 1 mg/mL. A scan rate of 60°C per hour was used with temperatures ranging from 20°C to 100°C. Results were analyzed with Origin 7.0 Microcal plug-in software (Malvern Instruments). Concentration-normalized data were fitted to a non–2-state model with 2 peaks to calculate melting temperature and calorimetric enthalpy of the individual domains after subtracting the heat capacity of the reference buffer.

### HDX-MS analysis

HDX-MS analysis was performed using Waters HDX Technology (Waters, Milford, MA). Both IEC pre-peak and main peak fractions were concentrated to 4 mg/mL prior to the HDX experiment. Samples containing 2 μL each were diluted 10-fold with either H_2_O buffer (25 mM sodium phosphate, pH 7.4) or D_2_O buffer (25 mM sodium phosphate, pD 7.4) at 25°C. After incubation for 0 (undeuterated), 0.5, 1, 5, 10, 30, 60, and 120 minutes, the labeled samples were quenched by adding 20 μL of ice-cold quench buffer (7.2 M guanidine, 500 mM Tris(2-carboxyethyl) phosphine hydrochloride, pH 2.4) and were then further diluted with 120 μL of ice-cold 0.1% formic acid solution. Immediately, 50 μL of diluted samples was digested by an Enzymate Pepsin Column (5 μM, 2.1 × 30 mm; Waters) at 20°C. The peptic fragments were collected with an Acquity BEH C18 VanGuard Pre-column (2.1 × 5 mm; Waters) and eluted into an Acquity BEH C18 column (1.7 μm, 1.0 × 100 mm; Waters) at 0°C. Peptides separated on the column were analyzed with a Xevo G2-XS QToF mass spectrometer (Waters). Data analysis was performed as previously described (23), and 3-dimensional structures of PE38 were drawn based on the structure 1IKQ from the Protein Data Bank.

### CD22 binding assay

A surface plasmon resonance–based characterization assay was used to measure the kinetic binding affinity (equilibrium dissociation constant [K_D_]) of moxetumomab pasudotox for CD22. An *anti-his* tagged antibody was immobilized to 2 flow cells of a sensor chip, using a standard amine coupling procedure. A reference surface was created by omitting the capture of CD22 on 1 flow cell. A dilution series of moxetumomab pasudotox was prepared and then injected in a serial-flow fashion across the reference surface and the CD22 surface, followed by regeneration of both flow cells. The data were reference flow cell subtracted and buffer blanked and then fit to a 1:1 binding model, using Biacore Evaluation software (Precision Antibody, Columbia, MD). This model yields an association rate constant (k_a_) and a dissociation rate constant (k_d_). The K_D_ for the interaction between moxetumomab pasudotox and CD22 was determined by the equation K_D_ = k_d_/k_a_.

### Internalization assay

Kinetic imaging analysis was used to determine whether moxetumomab pasudotox IEC pre-peak material was internalized into B-lymphoma cells. Moxetumomab pasudotox was conjugated with Alexa Fluor 647 (Thermo Fisher Scientific, Waltham, MA) and was bound to Daudi cells. The internalization of moxetumomab pasudotox was monitored with confocal fluorescence microscopy. Quantitative assessment of kinetic images of the cell membrane and cytoplasm over time demonstrated the internalization of moxetumomab pasudotox into the cytoplasm.

### ADP ribosylation assay

An ADP ribosylation assay was used to evaluate the ADP ribosylation activity of moxetumomab pasudotox in vitro. Moxetumomab pasudotox reference standard and test samples were serially diluted in ultrapure nuclease-free water and incubated with an ADP ribosylation assay system containing eukaryotic eEF2, NAD+, luciferase mRNA, complete amino acids mixture, RNase inhibitor, and wheat germ extract in a 96-well plate for 2 hours at ambient temperature. After incubation, an equal volume of Steady Glo Luciferase assay buffer and substrate (Promega, Madison, WI) were added to each well and the plate was incubated at ambient temperature with gentle shaking for 30 minutes before the luminescence was read on a plate reader. The amount of luminescence detected in each well was directly proportional to the amount of translated luciferase protein, which was inversely proportional to the amount of ADP ribosylation of eEF2 by moxetumomab pasudotox. The test sample was considered to be active if there was similar inhibition of luciferase mRNA translation as in the moxetumomab pasudotox reference standard. If there was no inhibition of luciferase mRNA translation, then the sample was considered to be inactive.

## RESULTS AND DISCUSSION

### Separation and identification of Asn-358 deamidated variant

As shown in Figure 1a, IEC analysis of moxetumomab pasudotox starting material led to a 2-peak profile, indicating the presence of 2 charge variants. The elution time of these variants (10.0 min and 12.5 min) were consistent with those in previous studies (21, 30), so these two IEC peaks were termed “pre-peak” and “main peak.” As shown in Figure 1b and 1c, successful enrichment of two variants was confirmed, and subsequent peptide mapping analysis revealed that the pre-peak fraction was mainly Asn-358–deamidated moxetumomab pasudotox (97.5%), whereas the main peak fraction contained only 9.6% deamidated Asn-358.

**Figure 1.**
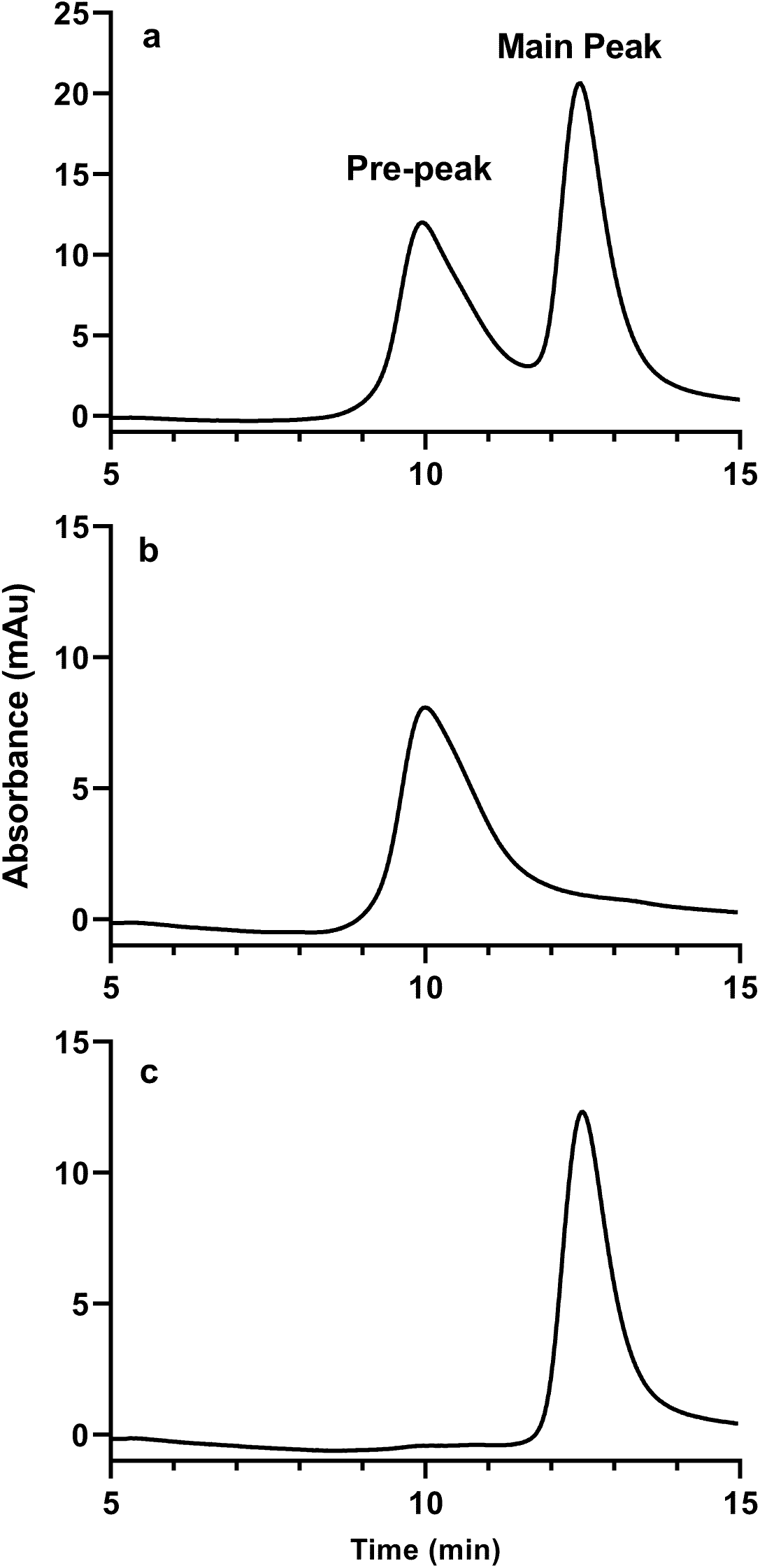
IEC fractionation of moxetumomab pasudotox. (a) IEC profile of starting material; (b) IEC profile of the pre-peak fraction; (c) IEC profile of the main peak fraction.

### DSC study

As a technology that can monitor the thermal unfolding transitions of a protein, DSC provided the first insight on the structural impact of Asn-358 deamidation. As shown in Figure 2 and Table 1, DSC measurements of both the pre-peak and the main peak fractions showed 2 distinct transitions, the first at ~44 °C and the second at ~78 °C. Native trypsin digestion of moxetumomab pasudotox was used to assign the first peak to unfolding of the PE38 domains and the second peak to the Fv domain (see Supporting Information), which is consistent with previous findings (32). Comparison of the two DSC profiles revealed that both fractions shared similar melting transition points for the Fv domain, but the PE38 domain of the pre-peak fraction showed a decrease of 1.3°C in melting temperature and a 15-kcal loss in enthalpy compared with the main peak fraction. These results suggest that Asn-358 deamidation had no effect on Fv thermal stability but slightly destabilized the PE38 region, indicating the possible conformation change in PE38.

**Figure 2.**
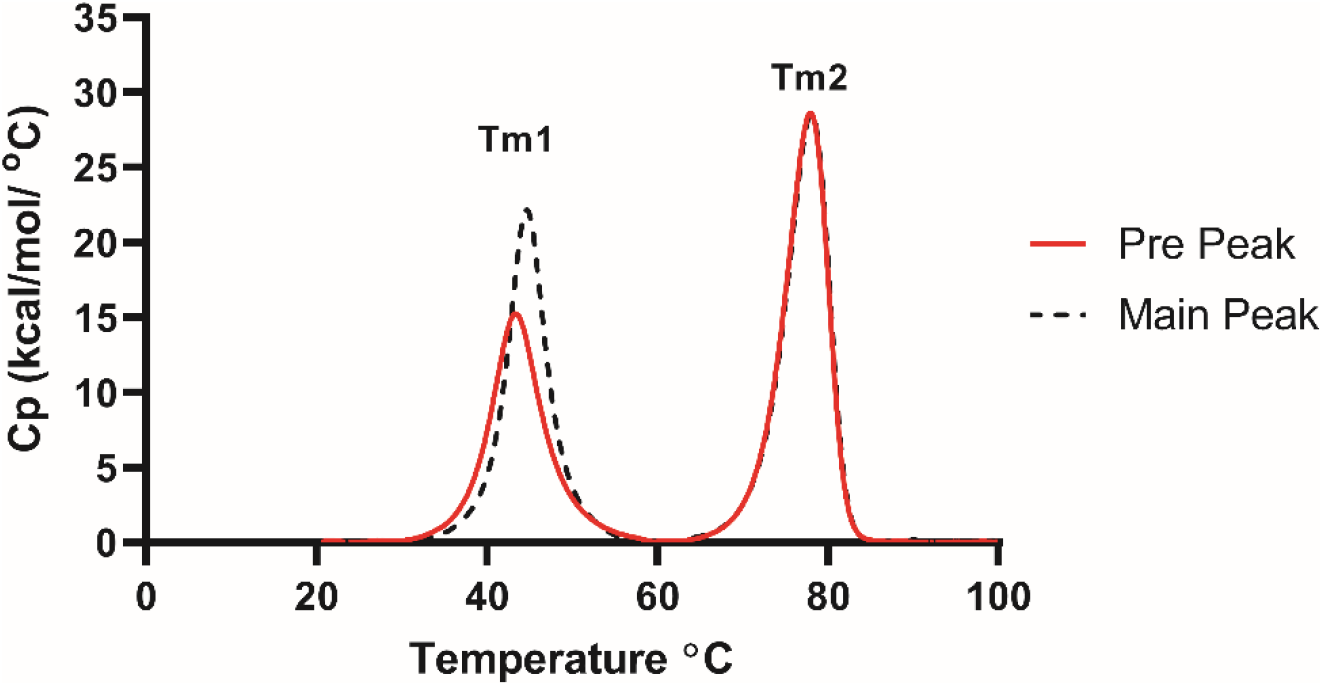
DSC thermograms of IEC pre-peak fraction (red solid line) and main peak fraction (black dash line).

**Table 1.** DSC melting temperature and enthalpy of the pre-peak and main peak fractions Tm = melting temperature.

| Sample | First transition |  | Second transition |  |
| --- | --- | --- | --- | --- |
|  | T<sub>m</sub> (°C) | Enthalpy (kcal) | T<sub>m</sub> (°C) | Enthalpy (kcal) |
| Pre-peak | 43.5 | 122 | 77.5 | 186 |
| Main peak | 44.8 | 137 | 77.6 | 186 |

T<sub>m</sub> = melting temperature.

### HDX-MS analysis

To further localize the conformational change caused by Asn-358 deamidation, HDX-MS was employed to analyze the two IEC fractions. A total of 182 peptic peptides were identified during the HDX data processing, and the standard deviation for each exchange time point was determined to be ±0.3 daltons (Da). Among all identified peptides, 14 were from the variable domain of light chain (LC), resulting in 81% of the LC sequence coverage. As shown in Figure 3a, both fractions had superimposable exchange kinetic curves for the peptide LC 37-47, indicating no conformational difference in this peptide region. Similar results were also observed for all the 13 other LC peptides. To gain a global view of the LC, the differences in deuterium incorporation between the 2 fractions were calculated for all the LC peptides (Figure 3b). It is clear that all these differences were smaller than 0.3 Da and were insignificant, so it was concluded that there was no structural difference between the two IEC fractions in the LC, which is consistent with DSC melting temperature for the second thermal transition.

**Figure 3.**
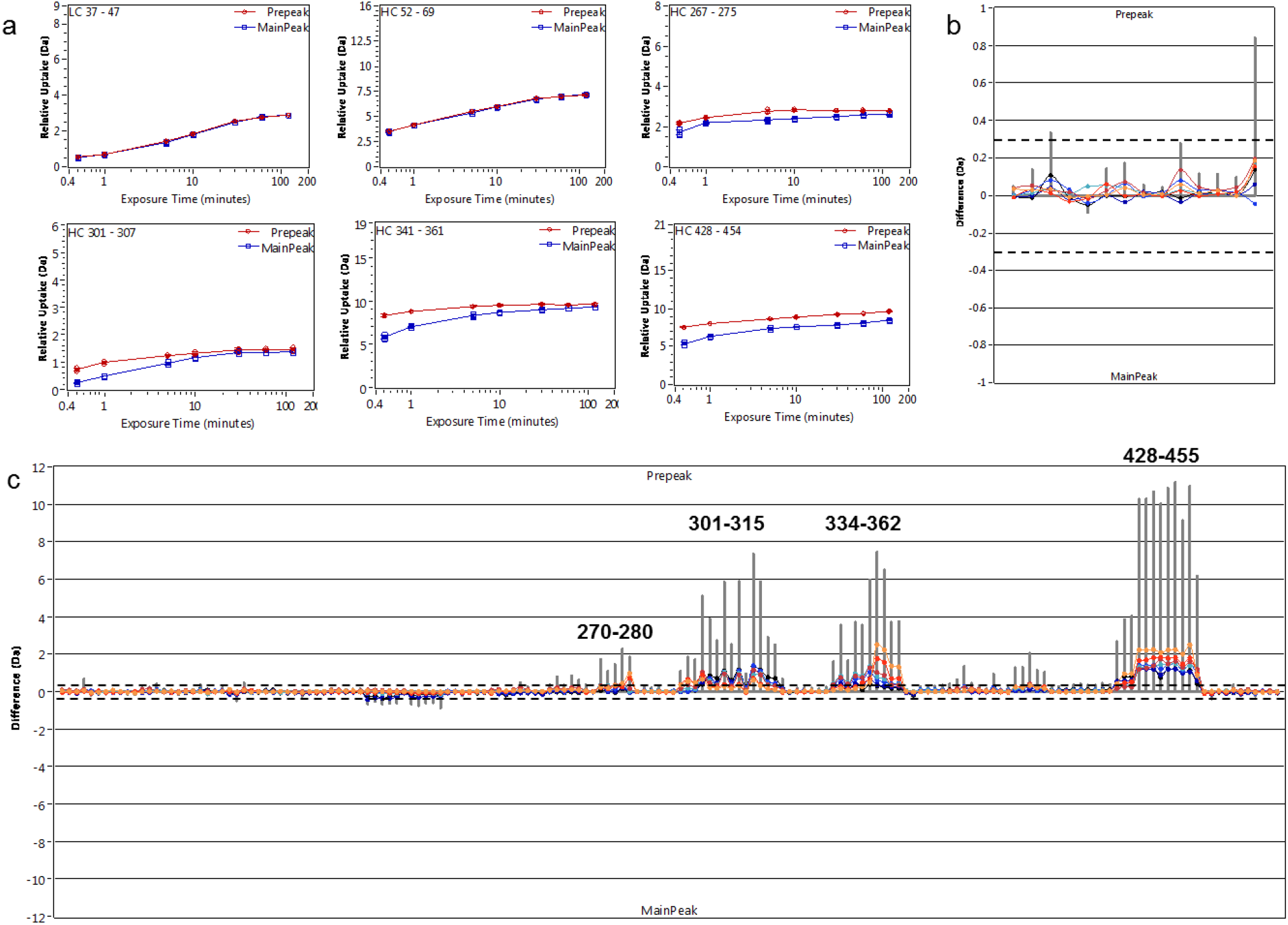
HDX analysis of pre-peak and main peak fractions. (a) Deuterium uptake kinetic curves of selected peptides from the pre-peak fraction (red) and the main peak fraction (blue). Difference charts of (b) light chain and (c) heavy chain summarize the difference in deuterium uptake between the 2 fractions for all 7 exchange time points: 0.5 minute (orange), 1 minute (red), 5 minutes (dark red), 10 minutes (cyan), 30 minutes (blue), 60 minutes (dark blue), and 120 minutes (black). The gray vertical bars represent the summed difference from all time points.

When the rest 168 peptides were identified to provide 96% sequence coverage of the VH and PE38 domains, which are also refer as heavy chain (HC), their HDX-MS data were more complex than those from the LC. First, consistent with the LC, all peptides located in the Fv region (residues HC1–129) showed no difference between the 2 fractions; exchange kinetic curves of peptides HC52–69 are shown as an example in Figure 3a. Second, most peptides in the PE38 region (residues HC130–476) showed no or only minor differences (≤0.3 Da) in deuterium uptake between the pre-peak and the main peak fractions, suggesting that no significant conformational difference was caused by Asn-358 deamidation in most of the PE38 region. Finally, obvious differences were observed for peptides associated with four regions (HC270–280, HC301–315, HC334–362, and HC428–455). As shown in Figure 3a, the peptides in regions HC267–275, HC301–307, HC341–361, and HC428–454 all exhibited increased deuterium uptake in the pre-peak fraction compared with the main peak fraction, and all other peptides related to these regions showed similar differences (Figure 3c). The differences at many exchange time points were greater than 0.3 Da, and a few were as high as 2.5 Da. These results indicated that some local conformational changes led to less protection against deuterium exchange in these regions.

One limitation of the differences shown in Figure 3c is that the peptide length was not considered, and sometimes longer peptides show more significant differences than short ones because they are able to accumulate more backbone amide deuterium, also because back-exchange can easily overwhelm the exchange on shorter peptides. To take the peptide length into account, the differences in deuterium uptake by PE38 peptides between the 2 fractions were divided by the total number of the exchangeable amide hydrogens in each peptide, depicted as colors in the heat map (Figure 4). This heat map clearly highlights the four PE38 regions in which the pre-peak fraction showed significantly higher deuterium uptake than the main peak fraction. Overall, the HDX data suggest that Asn-358 deamidation had no impact on the Fv region but did change local conformation in four regions in PE38. Considering that these regions account for only ~24% of total PE38 sequence, these data are consistent with the 1.3°C difference in the DSC melting temperature of the first transition.

**Figure 4.**
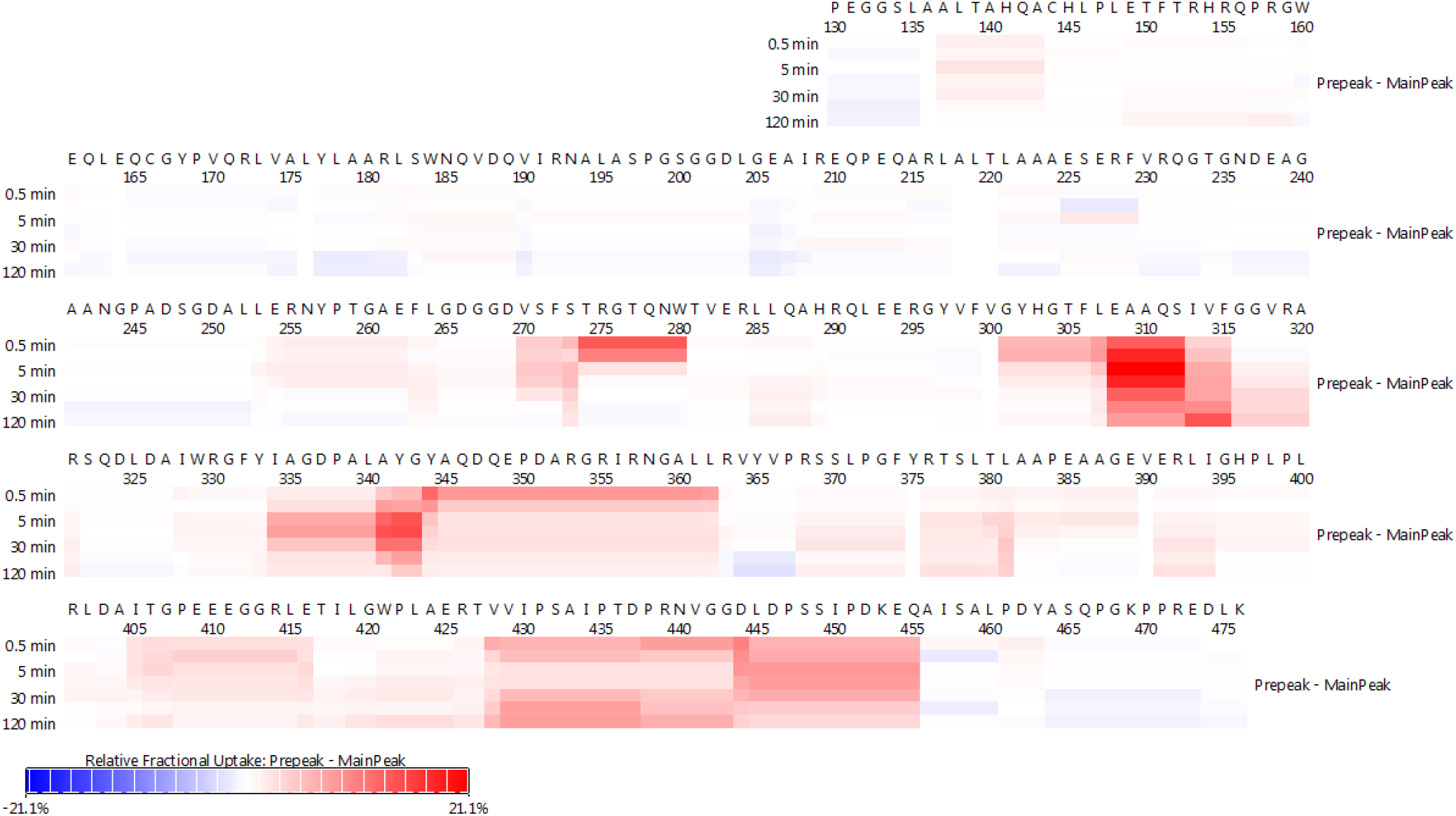
HDX heat map of PE38. Relative deuterium uptake differences between pre-peak and main peak fractions are shown in color along the sequence for all 7 exchange time points: 0.5, 1, 5, 10, 30, 60, and 120 minutes.

In addition, careful examination of the sequence revealed that these HDX-defined regions contained a total of 5 arginine residues. Because the HDX data suggested that these regions gained more solvent accessibility due to the deamidation of Asn-358, the deamidated moxetumomab pasudotox molecules might expose more positive charges that overcome 1 negative charge from Asn-358 deamidation; this would explain why the deamidated species eluted before the unmodified ones on a positively charged anion exchange column.

### Plausible mechanism for biological activity loss

As a truncated form of PE, PE38 contains part of the expression improvement domain (domain Ib), the translocation domain (domain II), and the ADP ribosyltransferase enzyme domain (domain III). The finding that all 4 HDX-defined regions were located in domain III suggested that the deamidation of Asn-358 (Asn-495 in PE) led to the misfolding of the catalytic domain and reduction of the biological activity of moxetumomab pasudotox. In a previous structural study of the complex eEF2-PE, eEF2 mainly bound to 486-493, 576-580, and Arg-412 of PE (33). When the HDX heat map and these eEF2 binding interfaces were demonstrated on the crystal structure of PE in our study (Figure 5), it became obvious that the eEF2 binding interface completely overlapped with the 4 HDX-defined regions 407–417, 438–452, 471–499, and 565–592 in PE (see Supplementary Information for sequence alignment of PE and PE38). Furthermore, 5 of 9 residues that constructed the NAD binding interface were also located in these HDX-defined regions (Figure 5b) (34). All these results suggest that the conformational changes resulting from Asn-358 deamidation disrupted binding of moxetumomab pasudotox to eEF2 and NAD, which led to loss of PE38-mediated cell killing biological activity.

**Figure 5.**
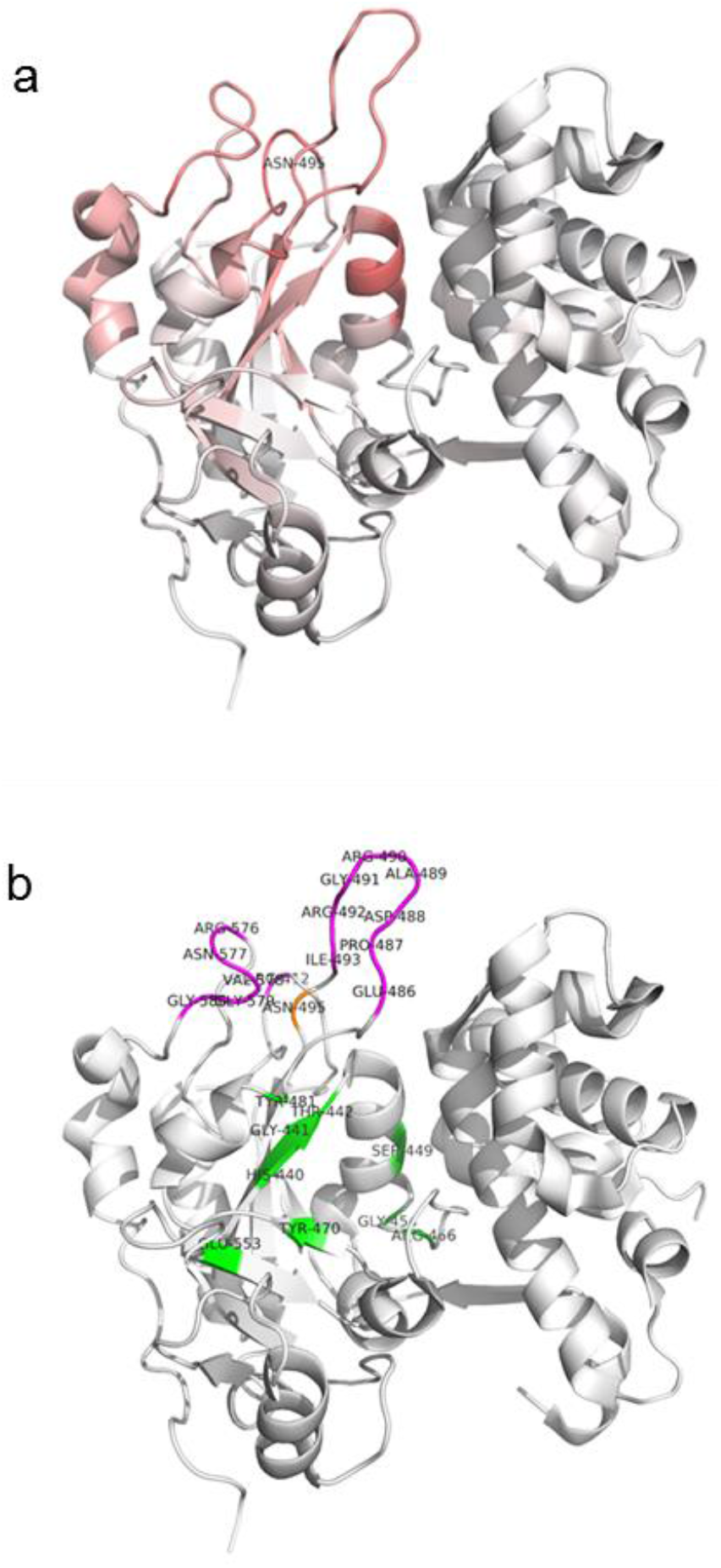
Crystalline structure of PE (PDB 1IKQ). (a) HDX heat map of 0.5-minute exchange time point; (b) binding interface of eEF2 (magenta) (33) and NAD (green) (34). The deamidation site is shown in orange. All residues are numbered according to PE.

This proposed mechanism is supported by observations from further biological characterization of the IEC pre-peak fraction as summarized in Table 2. The target binding results indicate that the IEC pre-peak material is capable of binding to the soluble CD22 with similar affinity as main peak, indicating that deamidation of PE38 at residue Asn-358 does not impact binding to the CD22. Kinetic imaging analysis was used to determine whether moxetumomab pasudotox the IEC pre-peak material is internalized into B-lymphoma cells. Quantitative assessment of kinetic images of the cell membrane and cytoplasm over time demonstrated the internalization of moxetumomab pasudotox into the cytoplasm. The IEC pre-peak material was internalized into the cells with a similar kinetics as moxetumomab pasudotox starting material (Supplementary figure S5). The ADP-ribosylation activity of moxetumomab pasudotox was assessed using an in vitro luciferase translation assay. No inhibition of the luciferase mRNA translation was observed for IEC pre-peak material, while the moxetumomab pasudotox starting material inhibited the luciferase mRNA translation in a dose dependent manner as shown in Figure 6. These results indicate that the IEC pre-peak material shows dramatically decreased activity to induce the ADP-ribosylation of eEF2, as indicated by the changes on the catalyst domain of PE38, and therefore caused the loss of cell-killing activity.

**Figure 6.**
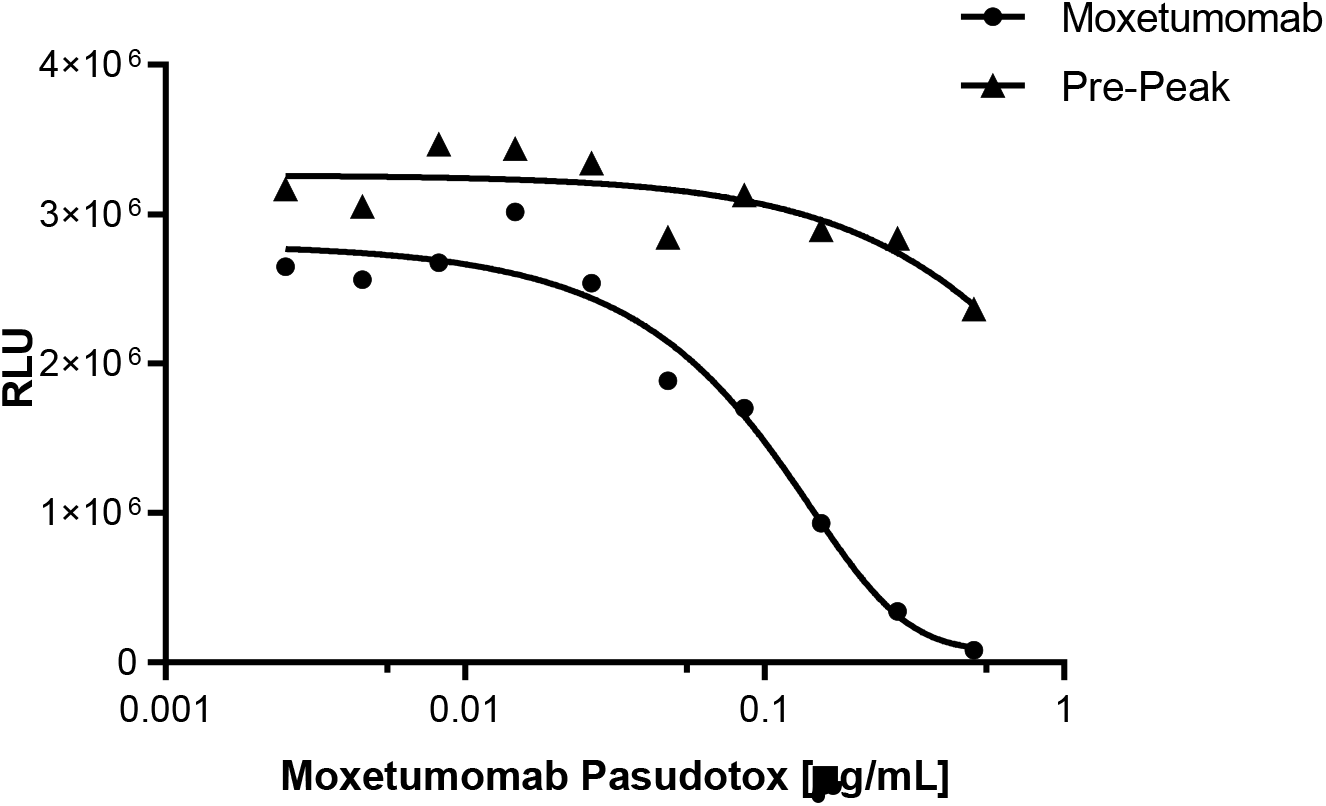
Dose-responsive curves of ADP ribosylation activity of IEC Pre-Peak (triangle) and starting moxetumomab pasudotox (circle).

**Table 2.** Summary of IEC pre-peak biological characterization

| Method | Purpose | Results |
| --- | --- | --- |
| CD22 binding assay | Measure binding affinity to CD22 | $k_D=1.39$ nM for Pre-peak<br>$k_D=1.38$ nM for main peak |
| Internalization assay | Measure cellular internalization | Internalization observed |
| ADP-ribosylation assay | Measure ADP ribosylation activity | Inactive |
| Apoptosis bioassay | Measure cell killing activity (relative potency) | Inactive |

The correlation of IEC pre-peak levels and relative potency by apoptosis bioassay were linearly plotted as described in a prior study (30), but the ratio of percent increase in deamidation to the percent decrease in relative potency was not 1:1. A spiking study was therefore carried out in which various ratios of purified IEC pre-peak to main peak were tested for relative potency. The correlation between IEC pre-peak to potency was nonlinear (Figure 7), indicating that the IEC pre-peak material was not only inactive but also appeared to competitively inhibit the biological activity of moxetumomab pasudotox. The relationship between the IEC pre-peak and relative potency can been expressed as:

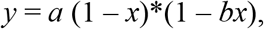

where *x* is percent pre-peak, *y* is percent relative potency, *a* is the relative potency of the pure main peak (percent pre-peak is 0), and *b* is the factor of competitive inhibition. In this case, *b* is approximately 0.55. With an understanding of the correlation between pre-peak and relative potency, an appropriate specification can be set for the pre-peak to ensure drug efficacy and safety.

**Figure 7.**
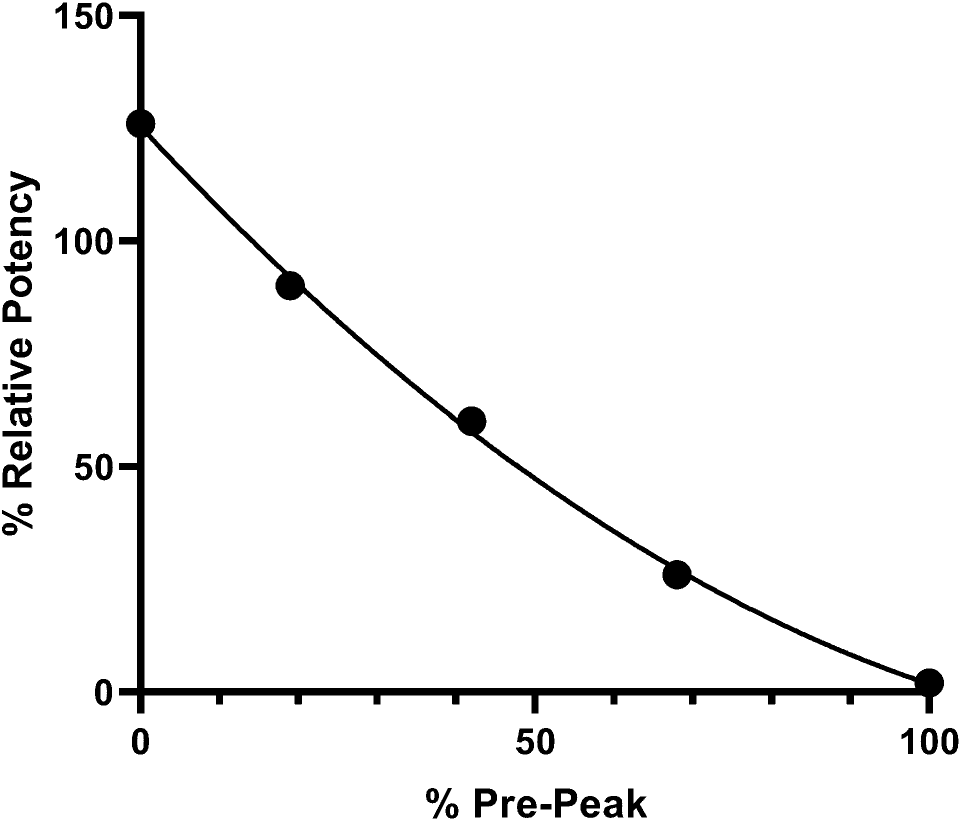
Correlation of IEC pre-peak ratio to relative potency.

Interestingly, the Asn-358 deamidation level did not change during the forced degradation study at high pH. The crystalline structure of PE indicates that Asn-358 (Asn-495 in PE) is buried, so Asn-358 deamidation can occur only during the refolding step, during which the molecule is exposed to high pH and denaturing conditions. The deamidated molecule folded incorrectly. With a clear understanding of the structural impact of the deamidation, process control can be achieved by optimizing the refolding conditions and the purification process to consistently produce high-quality product.

## CONCLUSIONS

In this study, we report a conformational change caused by single asparagine deamidation based on DSC and HDX MS data. It explained how Asn-358 deamidation impacted the biological activity of moxetumomab pasudotox and why the deamidated variant eluted as an early peak in anion exchange chromatography. Importantly, our results indicate that residue Asn-358 is enveloped once the protein is folded and thus will not change during storage conditions. With clear understanding of the structure-function relationship, the level of Asn-358 deamidation variant is minimized and controlled consistently in the current process for producing clinical and commercial supplies.

## AUTHOR INFORMATION

### Author Contributions

X.L. planned and organized all the experiments, conducted the HDX MS study and data processing, and wrote most of the manuscript. S.L. contributed to interpretation of biological data and contributed to the writing of the manuscript. N.D.M. performed the IEC fractionation and contributed to the writing of the manuscript. A.P. performed DSC and contributed to the writing of the manuscript. M.P. performed ADP ribosylation assay. J.D. updated the modeling and contributed to the writing of the manuscript. X.W. contributed to the writing, reviewing, and editing of the manuscript. J.W. performed peptide mapping analysis, summarized the data and conclusions, and contributed to the writing of the manuscript. All authors have approved the final version of the manuscript.

## Acknowledgments

All authors are either employee of AstraZeneca or was employee of AstraZeneca when the work was done. The authors acknowledge Timothy Pabst and Thomas Linke for providing the Moxetumomab Pasudotox material and LeeAnn Machiesky, Erika Farmer for testing biological activities, and Irma Vainshtein for internalization assay. We would like to thank Ben Niu and Xiaoyu Chen for review and edit. Editorial support was provided by Deborah Shuman (all are employee of AstraZeneca, Gaithersburg, MD, USA).

